# Deciphering tumour microenvironment and elucidating the origin of cancer cells in ovarian clear cell carcinoma

**DOI:** 10.1101/2024.08.06.606821

**Authors:** Uma S Kamaraj, Pradeep Gautam, Terence Cheng, Tham Su Chin, Sun Kuie Tay, Tew Hong Ho, Ravichandran Nadarajah, Ronald Chin Hong Goh, Shing Lih Wong, Sangeeta Mantoo, Inny Busmanis, Hu Li, Minh TN Le, Qi-Jing Li, Elaine Hsuen Lim, Yuin-Han Loh

**Affiliations:** Institute of Molecular and Cell Biology (IMCB), Agency for Science, Technology and Research (A*STAR), 61 Biopolis Drive, Proteos, Singapore 138673, Republic of Singapore; Department of Obstetrics & Gynaecology, Singapore General Hospital, Outram Road, Singapore 169608; Department of Anatomical Pathology, Singapore General Hospital, Academia, College Road, Singapore 169856; Center for Individualized Medicine, Department of Molecular Pharmacology and Experimental Therapeutics, Mayo Clinic, Rochester, MN 55905, USA; Yong Loo Lin School of Medicine, National University of Singapore, Singapore; Division of Medical Oncology, National Cancer Centre Singapore, 30 Hospital Boulevard, Singapore 168583; Department of Physiology, NUS Yong Loo Lin School of Medicine, 2 Medical Drive, MD9, Singapore, Singapore; Department of Biological Sciences, National University of Singapore, Singapore, Singapore; NUS Graduate School’s Integrative Sciences and Engineering Programme, National University of Singapore, 28 Medical Drive, Singapore, Singapore

## Abstract

Ovarian clear cell carcinoma (CCC) has an East Asian preponderance. It is associated with endometriosis, a benign condition where endometrial (inner lining of the uterus) tissue is found outside the uterus and on the peritoneal surface, in the abdominal or pelvic space. CCC is relatively more resistant to conventional chemotherapy compared to other ovarian cancer subtypes and is associated with a poorer prognosis. In this study, we recruited and obtained tumour tissues from seven patients across the four stages of CCC. The tumour and the tumour microenvironment (TME) from 7 CCC patients spanning clinical stages 1-4 were transcriptionally profiled using high-resolution scRNA-seq to gain insight into CCC’s biological mechanisms. Firstly, we built a scRNA-seq resource for the CCC tumour microenvironment (TME). Secondly, we identified the different cell type proportions and found high levels of immune infiltration in CCC. Thirdly, since CCC is associated with endometriosis, we compared CCC with two publicly available endometriosis scRNA-seq datasets. The CCC malignant cells showed similarities with glandular secretory and ciliated epithelial cells found in endometriosis. Finally, we determined the differences in cell-cell communication between various cell types present in CCC TME and endometriosis conditions to gain insights into the transformations in CCC.

## Introduction

Ovarian cancer (OC) is a lethal gynaecological cancer affecting women globally and is the 7^th^ most common cause of female cancer-associated death^1,2^. The World Health Organisation’s (WHO) International Agency for Research on Cancer (IARC) estimates ovarian cancer to increase in incidence and mortality in the next two decades^3^. The high mortality rates associated with OC are primarily due to the advanced disease stage at presentation and the lack of effective screening tools. It is categorised into four subtypes: serous, endometroid, clear cell, and mucinous^4^. Ovarian clear cell carcinoma (CCC) exhibits different prevalence rates across various regions, representing 5-10% of cases in North America but ranging from 13-25% in East Asia^5,6^. The greater East Asian preponderance is mirrored by the higher incidence of endometriosis in East Asia^7–10^, CCC being the subtype more highly correlated with endometriosis^11–16^.

Endometriosis is an inflammatory, non-malignant lesion affecting 10% of women of reproductive age worldwide^17^, causing pelvic pain and infertility. Sampson’s theory of retrograde menstruation^18^ postulated that endometrial cells moved from the inner lining of the uterus, through the Fallopian tubes, and into the abdominopelvic cavity. Women with endometriosis are three times more likely to develop CCC^16^. The pattern of distribution of endometriotic lesions in the abdominal or pelvic space is similar to ovarian cancer metastasis. Oncogenic mutations in *ARID1A*, *PIK3CA*, *KRAS* and *PPP2R1A*^19^ have been described in endometriotic tissue. *PIK3CA* mutation has been described as an early event in CCC^20^. Altogether, these data suggest that endometriosis is less-than-benign, prompting an enduring clinical question as to whether endometriosis is a precursor or facilitator of CCC.

Our study compares scRNA-seq data on CCC with endometriosis to identify the precursor cells for CCC. A recent study identified two expression subtypes in CCC: epithelial-like (associated with earlier stages) and mesenchymal-like (associated with later stages)^21^. Recently, single-cell RNA sequencing (scRNA-seq) technology^22,23^ has been applied to decipher normal and tumour conditions. Two recent cohort studies have used scRNA-seq to describe endometriosis^24,25^. However, there is currently a lack of high-resolution scRNA-seq data on CCC. To fill this gap, we present scRNA-seq profiling of CCC patients in our cohort, which includes various clinical stages of CCC. We used previously published studies on endometriosis to understand the differences and associations between CCC and endometriosis and, as well as to elucidate the origin of ovarian CCC cells. Analysing the differential genes expressed in heterogenous cell types present in CCC compared across the controls, we sought to acquire molecular insights that might clarify the role of endometriosis in CCC. Further, we provide insights into the critical TME cellular communications, thereby identifying molecules and pathways potentially targetable by existing drugs or further therapeutic development.

## Results

### Cellular composition in CCC tumour microenvironment

The samples of surgically resected ovarian CCC tissue from seven patients of different disease stages, including FIGO stages 1A, 2A, 2B, 3A and 4 were collected (Figure 1a, Table S1). The H&E staining of the CCC samples showed that these samples have abnormal morphologies with multilobed nuclei (Figure S1a). Of these, three patients were in stage 1 with cancer in a single ovary, two patients were in stage 2 where cancer has spread to the bladder or bowel, one in stage 3 where cancer spread into the ovary and one in stage 4 where cancer has metastasised and spread to other organs. Additionally, three of these seven patients were diagnosed with endometriosis along with CCC and are referred to as CCC-MD (Figure 1a). To understand the cellular composition within the tumour sample, we performed high-resolution scRNA-seq on the seven patient tissues and integrated the cells across the patients (Figure 1b). Following quality control, we obtained 13,543 cells across these CCC samples to dissect the tumour microenvironment (TME) (Figure S1b-e).

**Figure 1:**
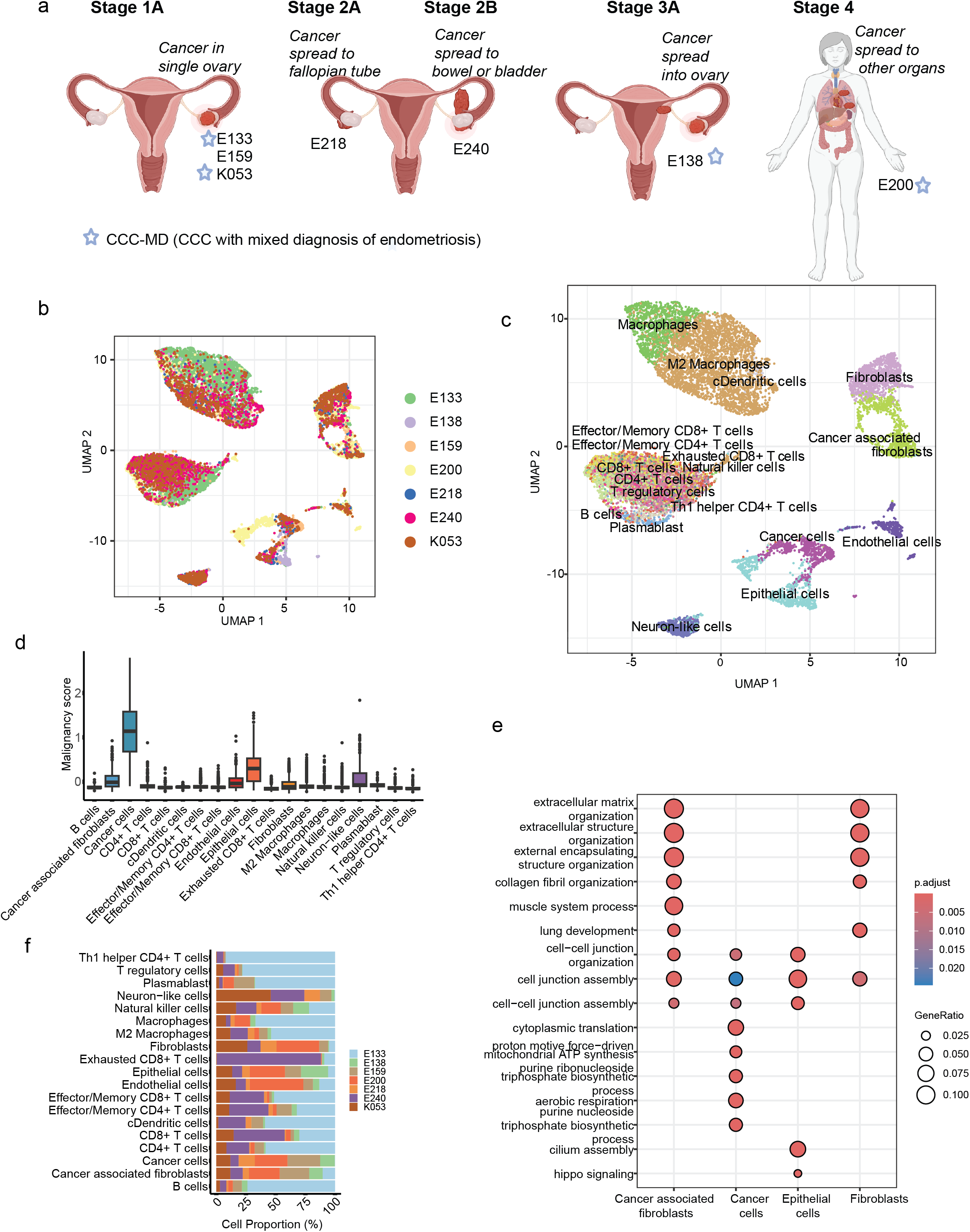
Overview of CCC samples used in this study and cellular composition of tumour microenvironment. a) The schematic shows the overview of the tumour samples from seven patients across various clinical stages of CCC. CCC refers to ovarian clear cell carcinoma, and CCC-MD refers to CCC with a mixed diagnosis of endometriosis. The schematic was created in BioRender.com. b) The integrated scRNA-seq data of the CCC tumour samples from patients were visualised using UMAP dimension reduction. c) The cell types identified in the samples were visualised using UMAP. d) The boxplot shows the malignancy score based on cancer-associated genes across all the cell types in CCC. e) The dot plot shows the gene ontology enrichment in selected epithelial cells, cancer cells, fibroblasts and cancer-associated fibroblasts across all the samples. f) The percentage of different cell types in the tumours of each of the patients.

To determine the cell types in the tumour samples, we used scATOMIC^26^, a model trained with the transcriptional profiles of the cell types annotated from publicly available scRNA-seq datasets on TMEs across 19 common cancers, including ovarian cancer. Using scATOMIC and manual curation, we annotated the different cell types in the CCC TME (Figure 1c). We observed 19 cell types in the CCC, including epithelial cells, endothelial cells, fibroblasts, neurons, and immune cells. The cells annotated as cancer cells have higher expressions of malignancy markers obtained from the CancerSCEM^27^ database (Figure 1d). Consistent with the previous studies, cells annotated as B cells express *CD79A*, *CD19*, and *MS4A1*^28^, exhausted T cells express *PDCD1*^29^, Treg cells express *FOXP3*^30^. Macrophages in CCC samples were separated into two groups, M2 macrophages and macrophages based on the high expression of *CD86* and low expression of *CCL5*^31^. Further, cancer-associated fibroblasts were annotated based on the expression of *ACTA2* and *TAGLN*^32^ (Figure S1f). The epithelial and cancer cells showed higher *KRT8*, *KRT18*, *KRT19* and *ANXA4* expression^27^.

Subsequently, differential gene ontology biological processes indicated that the cancer cells had enrichment for cell division and proliferation-related terms compared to the epithelial cells, and the cancer-associated fibroblasts showed enrichment of cell-cell assembly-related gene ontology biological processes (Figure 1e, S1g). *ARID1A* and *PIK3CA* are well-known oncogenes associated with CCC^33^, and here, we also found a higher mutation rate at these genes across the samples (Figure S2a-b). Based on cell population, a significantly higher proportion of immune cells were present in all the CCC tumours compared to epithelial or fibroblasts, indicating an immune infiltration at the tumour site (Figure 1f).

### Immune infiltration and cellular communication in TME

Henceforth, we sought to decipher the cell-cell communications within the TME of the collected patient tumour samples. The top signalling pathways active in the TME across all samples were pathways such as MIF, TGFb, PARs, SPP1 and CXCL (Figure S2c). The cancer cells in CCC predominately secreted ligands for pathways such as MIF and SPP1 (Figure 2a). The cancer cells and cancer-associated fibroblasts produced the MIF ligands, signalling to the immune cells such as macrophages and T cells (Figure 2b). Interestingly, the surrounding normal epithelial cells in CCC are not involved in *MIF*-*CD74* signalling and are cancer-specific. *MIF* cytokine is known to have a pro-tumour role with functions covering several hallmarks of cancer, including resisting cell death, inducing angiogenesis, promoting genome instability, tumour proliferation and acting as an immunosuppressor^34^. Similarly, the SPP1 pathway showed significant signalling interaction between cancer and macrophages, and *SPP1* is a poor prognostic marker for clear cell carcinoma^35,36^. Cancer-associated fibroblasts signal to the cancer cells for tumour growth and angiogenesis via pathways such as HGF, ANGPTL4, HBEGF and FGF (Figure 2d). Dysregulation of the *HGF*-*MET* axis is well-known in tumorigenesis and invasion^37^, and here in CCC, cancer-associated fibroblasts and macrophages signal the cancer cells (Figure 2e). Similarly, *PPIA* and *BSG* co-expression is significantly associated with poor prognosis in tumours^38^, and in CCC the signalling involves all the immune and cancer cells (Figure 2f).

**Figure 2:**
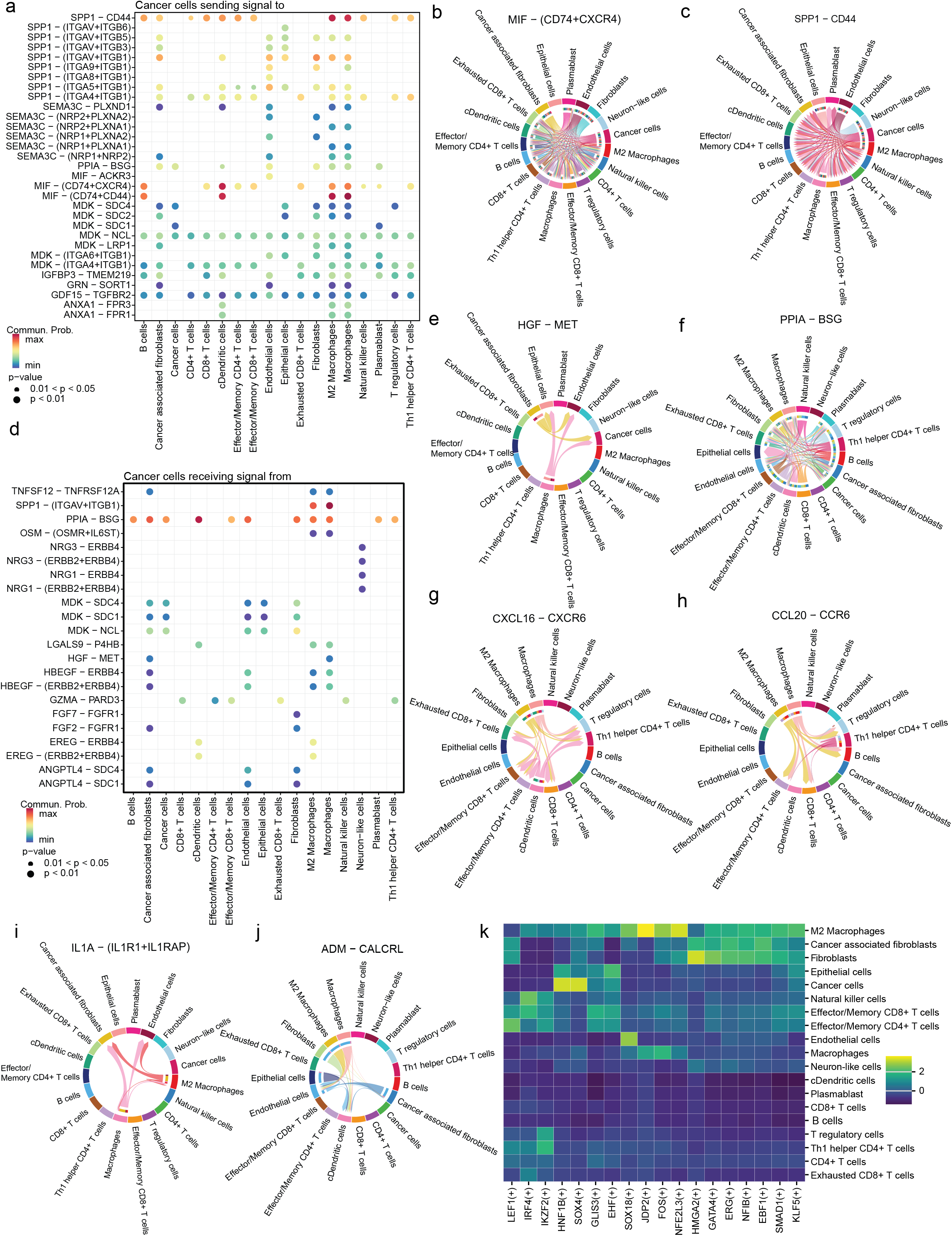
Cell-cell communication signalling in the tumour microenvironment: a) The dot plot shows the significant ligands the cancer cells produce targeting the receptors in the ligand-receptor (L-R) signalling. The chord plots show the ligand and receptor interaction among the cell types in the TME for the following selected signalling pathways: b) MIF pathway with MIF ligand and *CD74*-*CXCR4* receptor, c) SPP1 pathway with *SPP1* ligand and *CD44* receptor. The colour of the connections between cell types represents the ligands expressed from the cell type and the arrow points to the receptor of the cell types. d) The dot plot shows the significant receptors that are active in the cancer cells for the ligands produced by other cell types in the TME. The chord plots show the ligand and receptor interaction among the cell types in the TME for the following selected signalling pathways: e) HGF pathway with HGF ligand and MET receptor, f) CypA pathway with *PPIA* ligand and *BSG* receptor, g) CXCL pathway with *CXCL16* ligand and *CXCR6* receptor, h) CCL pathway with *CCL20* ligand and *CCR7* receptor, i) IL1 pathway with *IL1A* ligand and *IL1R1*-*IL1RAP* receptor and j) CALCR pathway with *ADM* ligand and *CALCRL* receptor. k) The heatmap shows the enrichment of TFs regulons with regulon specificity scores in the cell types found in TME.

The other top chemokine signalling pathways in TME were CXCL and CCL signalling and macrophages show significant secretion of these chemokines (Figure S2d). For instance, *CXCL16* is a chemoattractant for the Treg cells to the tumour site and its pro-tumourigenic functions^39^, and here we show the macrophages and dendritic cells recruits Treg, Th 1 helper CD4+ T and other CD8+ T cells (Figure 2g). Recently, the *CCL20*-*CCR6* axis has been known to promote cancer progression by enabling cancer cell migration and proliferation via CCL20 signalling also known as macrophage inflammatory protein^40^ and in CCC macrophages recruit Treg, CD8+ T and effector CD4+ T cells (Figure 2h). Interleukins play a critical role in promoting cancer progression while essential for tumour-directed immune response^41^, the interleukins prominent in the CCC TME are *IL1*, *IL2*, *IL6*, *IL16* and *IL10* (Figure S2d). IL1 signalling promotes angiogenesis^42^, and macrophages direct signalling to endothelial cells to promote angiogenesis. Furthermore, *IL1* is shown in the conversion of macrophages towards the M2 phenotype^42^, and there is evident *IL1A* signalling between the two macrophage populations in TME (Figure 2i). Another pathway is the CALCR pathway with the *ADM*-*CALCRL* communication, the macrophages and cancer-associated fibroblasts signal the endothelial cells via ADM secretion, suggesting the promotion of angiogenesis (Figure 2j). *ADM* has been indicated in the upregulation of the *VEGF* pathway promoting angiogenesis in ovarian carcinoma^43^. This shows that cellular communication in the CCC TME promotes angiogenesis and has a pro-tumourigenic microenvironment in the patients.

### Master regulators governing CCC TME

The key regulatory transcription factors (TFs) for each cell population in the CCC TME were identified using SCENIC^44^. The critical master regulators of CCC cancer cells were *HNF1B* and *SOX4* (Figure 2k, S2e). *HNF1B* is a hepatocyte-specific TF expressed in several cancers, and the reduction of *HNF1B* levels in CCC cell lines has been shown to cause apoptosis^45^. Similarly, *SOX4* is upregulated in subtypes of cancerous cells of clear cell renal carcinoma^46^. *SOX4* plays a key role in maintaining the stemness of cancer cells^47^ and is a known master regulator of EMT pathways in breast cancer^48^. M2 macrophages in CCC showed high *NFE2L3* regulon activity and *NFE2L3* expression is a poor prognosis marker in colon, pancreatic and renal cancers^49–51^. A study has also shown that loss of *NFE2L3* protects against inflammation caused by colorectal cancer by modulating TME^52^. Interestingly, exhausted T cells had higher regulon activity of *IRF4* compared to CD4 and CD8 T cells (Figure 2k). *IRF4* has been shown to drive the exhaustion of CD8+T cells in chronic infection^53^. Overall thesignalling pathways and transcriptional regulators in CCC are pro-tumourgenic.

### Progression of CCC cancer cells

Next, we aimed to study the nature of CCC cancer cell progression across the four cancer stages. Interestingly, the more advanced stages of cancer stages 3 and 4 have reduced responses to chemokines and humoral immune responses based on differential gene ontology (Figure S3a). The chemokines are usually responsible for the recruitment of immune cells to the tumour site and humoral immunity comprising of B cells is important for tumour suppression. Here, the advanced stages of CCC seem to be immune-compromised. Further, cell trajectory analysis was performed to map the progression of cancer cells in CCC (Figure 3a, S3b-c). We observed that the stage 1 cancer cells are on one end of the spectrum, and metastasised stage 4 cancer cells are on the other, demonstrating the cancer cells’ overall progression. Next, the cell fate probabilities were determined to identify the gene expression changes during cancer stage progression (Figure S3c). The sequential genes upregulated during the progression from cancer stage 1 to 4 are *CALB1*, *TPD52L1* and *ITGB8* (Figure 3b). *CALB1* is an oncogene known in ovarian cancer and acts by inhibiting p53 pathway^54^. Similarly, *TPD52L1*^55^ *and ITGB8*^56^ are known to be in other cancer transitions. Additionally, in metastasised stage 4 cancer, ligands such as *GDNF* and *DKK1* are also responsible for tumour progression^57^, especially *DKK1* is known to promote tumour metastasis^58^.

**Figure 3:**
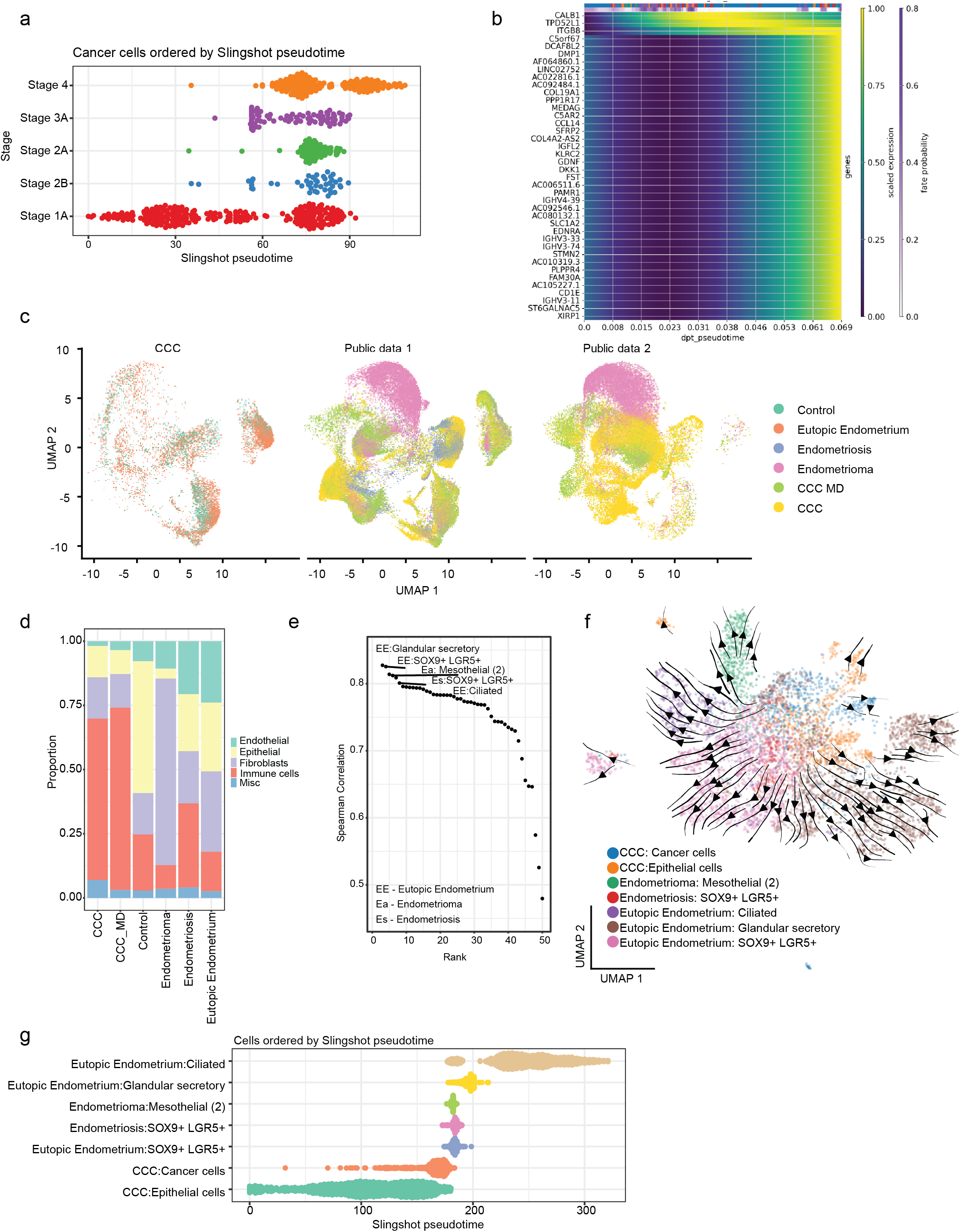
Deciphering the CCC cancer cells progression and origin of epithelial cells. a) The cancer cells are ordered by the stages of cancer based on slingshot pseudotime. b) The changes in gene expression across the diffusion pseudo-time from stage 1 to 4 cancer cells, along with the cell fate probability changes. c) UMAP visualisation of integration of CCC tumour samples in this study with healthy control, eutopic endometrium, endometrioma and endometriosis from two publicly available datasets. d) The proportion of cell groups present in each condition. e) The Spearman correlation of the transcriptional profiles between the cancer cells in CCC and the epithelial cells found in healthy controls and other endometrial conditions. The cell types with a Spearman correlation greater than 0.8 with CCC cancer cells are labelled. f) The UMAP visualisation of the CCC cancer cells and the correlated epithelial cell types. The projection lines and arrows represent the pseudotime. g) The correlated epithelial cells are ordered by slingshot pseudotime.

### Cellular composition of various endometrial conditions

Since CCC is associated with endometriosis, we compared CCC with healthy endometrium and endometriosis-related conditions using two publicly available scRNA-seq datasets^24,25^. Firstly, we combined the single cells in the two datasets and grouped them as “control” representing healthy endometrium, “eutopic endometrium” representing the healthy endometrium of donors with endometriosis, “endometriosis” representing the tissue with endometriosis and “endometrioma” which are cystic lesions that stem from endometriosis. The integrated scRNA-seq data from this study and public datasets comprising six conditions such as control, eutopic endometrium, endometriosis, endometrioma, CCC and CCC-MD (CCC with a mixed diagnosis of endometriosis) are shown via UMAP dimension reduction projection (Figure 3c).

To understand the composition of the tissue in different endometrial conditions, the proportion of cell types present in each condition was determined (Figure 3d, Figure S3d-g). The endometrial region has an abundance of immune cells, with the healthy control containing more than 20% immune cells and 50% epithelial cells. During the endometriosis condition, we can already observe an increase in immune and endothelial cells by more than 10% and a reduction in the proportion of epithelial cells. The population of immune cells further increases to 65% in cancerous conditions like the CCC and CCC-MD, suggesting an immune infiltration. Meanwhile, the endometrioma cyst is predominantly made of fibroblasts and fewer immune cells.

### Deciphering the origin of CCC cancer cells

Endometriosis is defined by the presence of endometrial tissue in regions other than the endometrium. As women with endometriosis are more likely to get CCC, we aimed to decipher whether endometriosis is a precursor or facilitator of CCC by comparing the cancer cells with the epithelial cells found in endometrium and other endometriosis conditions. The transcriptional similarity was detected by Spearman correlation between CCC cancer cells and epithelial cells found in other conditions (Figure 3e, Table S2). The transcriptionally closest epithelial cells were glandular secretory, SOX9+ LGR5+ epithelial cells and ciliated in the eutopic endometrium of donors with endometriosis. The glandular secretory and ciliated epithelial cells are found in the luteal phase, and SOX9+ LGR5+ epithelial cells are found in the follicular phase of the menstrual cycle^25^. These cell types are transcriptional-related closely based on UMAP projection and pseudotime trajectory (Figure 3f-e). Further, gene ontology analysis also reveals the similarity between the cell types (Figure S3h). The one key signalling pathway active in cancer cells and not in the other compared to epithelial cells was the CyP signalling, known in poor tumour prognosis^38^ (Figure S4a, Figure 2f). Interestingly, it was shown that glandular secretory epithelial cells have high expression of *ARID1A* and *KRAS*^25^ and this is correlated with ovarian clear cell carcinoma samples^59^. These findings suggest that the CCC cancer cells might have originated from the epithelial cells related to the endometriosis condition such as glandular/secretory, ciliated and SOX9+ LGR5+ epithelial cells.

### Cellular communication in the microenvironment of endometrial conditions and CCC

It is necessary to decipher the differences in cell-cell communication to understand the dynamics of different endometrial conditions and CCC. Here, we performed cell-cell communication analysis based on the transcriptional profiles of the ligands and receptors in cell types found in the microenvironment. The major signalling pathways specific to the cancer condition are VEGI, APRIL, IL10, and CD70 (Figure 4a), and these pathways mostly involve the signalling from immune cells. The macrophages recruit the T cells, including the Treg and Th1 helper cells via VEGI (TNFSF15) signalling (Figure 4b), and it has been shown that *TNFSF15* promotes macrophage polarisation toward M1 for tumour suppression^60^. On the other hand, *APRIL* (*TNFSF13*) is known to stimulate B cell growth (Figure 4c) and cancer cell growth^61^. Further, *IL10* an anti-inflammatory ligand, is secreted by M2 macrophages to Th1 helper T cells and dendritic cells (Figure 4d). Finally, dysregulation of the CD70-CD27 axis within TME has been associated with tumour progression and immunosuppression^62^, and we observed significant signalling between B cells, Tregs, and exhausted T cells (Figure 4e). The *OX40* and *CD137* show differential presence in CCC and CCC with mixed diagnosis of endometriosis. Interestingly, the CALCR and BTLA pathways are present in both endometriosis conditions and CCC but not in healthy tissue, indicating a dysregulation in the calcium pathway and immune checkpoint, respectively.

**Figure 4:**
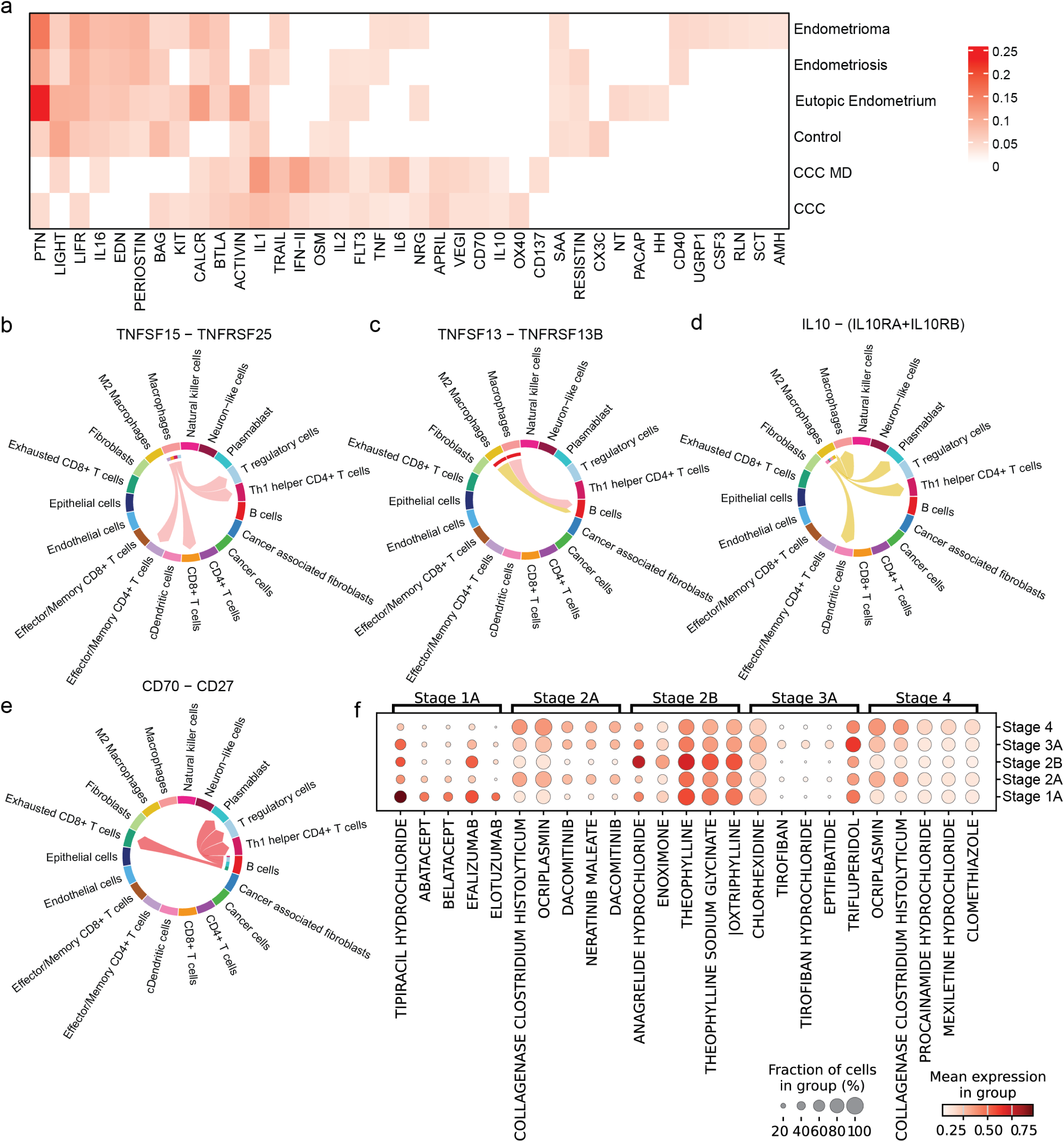
Comparison of cell-cell communication between conditions and prediction of potential drugs in CCC. a) The heatmap shows the distinct signalling pathways enriched across different conditions. b) The predicted drugs against the transcriptome profile of the TME across the stages of cancer. The chord plots show the ligand and receptor interaction among the cell types in the TME for the following selected signalling pathways in CCC for b) APRIL pathway with *TNFSF15* ligand and *TNFRSF25* receptor, c) VEGI pathway with *TNFSF13* ligand and *TNFRSF13B* receptor, d) IL10 pathway with *IL10* ligand and *IL10RA*-*IL10RB* receptor, e) CD70 pathway with *CD70* ligand and *CD27* receptor. The colour of the connections between cell types represents the ligands expressed from the cell type and the arrow points to the receptor of the cell types. f) The dot plot shows the predicted drugs targeting the transcriptional profile of the TME at different stages of cancer.

### Repurposing drugs targeting the TME

To predict the potential repurposing drugs for CCC, we aimed to predict the drugs based on the transcriptional profile of the TME. We stratified the patients based on the stages of cancer and used drug2cell^63^ to predict the drugs that could potentially target the TME (Figure 4f, S4b-c). The most significant drug predicted for Stage 1 TME is Tipiracil hydrochloride, which is a thymidine phosphorylase inhibitor used for the treatment of colorectal cancer^64^. The remaining top predicted drugs for Stage 1 were Abatacept and Belatacept targeting the T cells, Efalizumab targeting lymphocytes and Elotuzumab, which has an anti-tumour function by enhancing activation of natural killer cells based on the DrugBank database^65^. Some of the top predicted drugs for Stage 2 were Dacomitinib and Neratinib, and these drugs are used to treat non-small cell lung cancer and breast cancer, respectively. The drugs predicted for higher stages of CCC were drugs for preventing cardiovascular or asthma, suggesting the effects of medication the patients are already taking. Overall, the scRNA-seq profiling of CCC TME enables a detailed understanding of cell-cell communication and the possibility of identifying drugs for repurposing and working towards helping in personalised medicine.

## Discussion

This is the first study to profile single cells from clear cell carcinoma TME during different stages of cancer progression. We decipher the cellular communication in the TME and compare it with other endometrial-related conditions. We further show the potential of scRNA-seq to predict drugs for treatment. However, this study has some limitations, such as a smaller sample size and here we aim to use this as a discovery cohort to understand CCC progression. Another limitation is profiling was done in a single modality, therefore, as an extension, multimodal sequencing can be applied to comprehend CCC TME dynamics such as chromatin accessibility^66^, histone profiling^67^ and RNA modification^68^.

Across all donors, *MIF* and *SPP1* signalling was prominent in cancer cells while *SPP1* signalling was also active in M2 macrophages. The potential recipients of *SPP1* signalling included immune cells infiltrating tumours, cancer-associated fibroblasts, and regulatory T cells. Interestingly, *SPP1* signalling was high in CCC compared to endometrial conditions. A recent pan-cancer analysis using public data from TCGA showed that *SPP1* is overexpressed in most cancers^69^. *SPP1* has been shown to recruit macrophages to turn into tumour-associated macrophages in gliomas^70^. Anti-*SPP1* antibodies have been shown to suppress colon tumour growth in vivo^71^. *SPP1* inhibitors like parecoxib, brefelamide and simvastatin could be used as adjuvants in chemotherapy for the treatment of CCC.

Examination of the CCC TME revealed activation of several signalling pathways (HGF, ANGPTL4, HBEGF, and FGF) that promote tumour growth and angiogenesis. M2 macrophages displayed prominent *PPIA*-*BSG* signalling, previously linked to poor prognosis in various cancers. Infiltrating immune cells showed active *CXCL16* and CCL20 signalling, which aid cancer progression. Furthermore, TME recruitment of interleukins (*IL1*, *IL2*, *IL6*, *IL16*, and *IL10*) further promotes angiogenesis. Finally, *ADM* signalling, which upregulates VEGF signalling to promote blood vessel growth, was active in macrophages, CAFs, and endothelial cells.

Gene regulatory network analysis revealed distinct transcription factor activity in different cell types within the CCC tumour microenvironment. Exhausted T cells displayed the strongest regulon activity of *IRF4*, a transcription factor known to drive T-cell exhaustion during chronic inflammation. Blocking *IRF4* could be a potential therapeutic strategy to reinvigorate exhausted T cells and enhance anti-tumor immunity. Cancer cells, on the other hand, showed high expression of *HNF1B*, contributing to maintaining stemness in cancer cells, a characteristic essential for tumour growth. This finding suggests that *HNF1B* is another potential target for therapeutic intervention in CCC.

To understand the unique molecular signature of CCC, we compared it with endometriosis and healthy endometrium. As prior research indicates, women with endometriosis have a threefold increased risk of developing CCC. This comparison aimed to identify potential causative links by analysing cell-cell communication patterns within both conditions. We determined pathways specific to CCC, such as OX40 and CD137, by excluding signalling pathways common to endometriosis and healthy endometrium.

Interestingly, CCC cancer cells displayed transcriptional similarity to glandular secretory, ciliated, and SOX9+ LGR5+ epithelial found in endometriosis conditions. Furthermore, ARID1A, a well-established oncogene in CCC, is highly expressed in these same endometriosis cells^25^. This is particularly noteworthy as mutations in *ARID1A* are found in 57% of ovarian CCC cases^72^. Based on this shared expression of *ARID1A* and the observed transcriptional similarities, we postulate that ovarian CCC cells may originate from these specific epithelial cells present in the endometriosis condition.

Finally, we aim to find druggable targets during different stages of cancer progression. Tipiracil hydrochloride, which has been used for the treatment of colorectal cancer, was shown to be drug-specific for stage 1 of CCC. For Stage 2, Dacomitinib and Neratinib were predicted to counter CCC. These drugs are used to treat non-small cell lung cancer and breast cancer, respectively. Established based on this cohort, we show how we could determine potential candidates using scRNA-seq for patient-specific drug interventions in ovarian CCC.

## Supplementary Data

**Table S1:** Summary of the tumour samples used in this study

**Table S2:** Spearman correlation between cancer cells in CCC and the epithelial cells found in different endometrial conditions.

**Figure S1:** a) The H&E staining of the tumour tissue obtained from CCC patients. b) The bar plot shows the number of cells sequenced in the patient’s tumour samples. c) The number of unique genes detected across cells in the samples. d) UMAP visualisation of the cell-cycle phase of the patient cells. e) The percentage of mitochondrial genes detected in the samples. f) The dot plot shows the expression of marker genes among the identified cell types. g) The dot plot shows the gene ontology enrichment for immune cell types in the TME.

**Figure S2:** Mutation signature in a) ARID1A and b) PIK3CA genes with the SNP position and type represented on the protein structure across all the patients. c) The heatmap shows the top 20 signalling pathways enriched in the cancer TME across all the patients. d) The heatmap shows the signalling pathway patterns in each cell type based on the gene expression of outgoing ligands and incoming receptors. e) The heatmap shows the enrichment of TFs with regulon specificity scores in cancer cells across the patients.

**Figure S3:** a) Gene ontology biological processes enriched in cancer cells across the stages of CCC cancer. b) UMAP projection of cancer cells in all 4 stages. c) The UMAP visualisation of CCC cancer cells and the projection lines and arrows represent the diffusion pseudotime. The fate probabilities of diffusion pseudotime from cancer stage 1 to 4 cells. d) The cell proportion of annotated immune cells in different conditions. e) The cell proportion of annotated epithelial cells in different conditions. f) The cell proportion of annotated fibroblasts in different conditions. g) The cell proportion of annotated mural cells or neural-like cells in different conditions. h) Gene ontology biological processes that are enriched in selected epithelial cells which show a higher correlation to CCC cancer cells.

**Figure S4:** a) The heatmap shows the signalling pathways enriched in the epithelial cells that are transcriptionally closer to cancer cells in CCC. b) The heatmap shows the distinct signalling pathways enriched across different endometrial conditions. c) The dot plot shows the predicted drugs targeting the transcriptional profile of the cell types found in the TME across the CCC patients.

## Methods

### Sample acquisition

This study’s approval was obtained from the SingHealth Centralized Institutional Review Board, encompassing the National Cancer Centre Singapore, Singapore General Hospital, and Kandang Kerbau Women’s and Children’s Hospital (CIRB 2015/2595). Tissue samples from ovarian cancer patients with informed consent were obtained ex vivo and dissected freshly by the pathologist. Samples were collected in RPMI 1640 medium and delivered on ice to the laboratory within 1 hour of tissue collection.

### Processing the sample

Tissue samples of ovarian tumours were cut into ∼1-2 mm^3^ pieces using sterile scalpels and digested with collagenase type IV (STEMCELL Technologies) for 2 hours at 37°C on a rotator. Cell suspensions were filtered through a 70 mm cell strainer and centrifuged, and red blood cells (RBC) in the cell pellets were removed through RBC lysis. Cells were washed with PBS and snap frozen with liquid nitrogen and stored at -80°C until ready for single-cell analysis experimental use. The sample was dissociated, and the single-cell suspension was prepared by trypsin treatment for 2-3 hours at 37 C. The single-cell suspension was processed for single-cell analysis using 10× genomics single-cell RNAseq protocol. The single-cell library was sequenced using the Hiseq 4000 Illumina sequencer.

### scRNA-seq data processing

Cellranger v7.1 was used to map the 10X scRNA-seq data to GRCh38 v32 human reference genomes. The resulting count data was analysed using Seurat v4.3^73^. The cells were filtered out if the cells contained less than 500 genes or greater than 25% mitochondria reads. The cells with UMI counts less than 0.01% quantile and greater than 95% quantile across all cells were removed. The genes were filtered out if expressed in less than 10 cells. The genes with log10 of average UMI count across all cells greater than -2.5, genes with at least 2 UMIs and genes detected in at least 10 cells were used for downstream analysis. DoubletFinder v2.0.3^74^ was used to identify and remove doublets.

### Cell type annotation

Next, to annotate the cell types, we used scATOMIC^26^ a machine-learning cell annotation method trained using public pan-cancer scRNA-seq data. We additionally manually curated the annotation using well-known marker genes for each cell type.

### Integration of across samples and studies

The gene count for each tumour sample was normalised using SCTransform^75^ and dimension reduction was performed using PCA and UMAP. The seven samples were integrated using the Pearson residuals integration method from Seurat SCTransform integration, resulting in 13,543 cells.

We integrated publicly available endometriosis datasets^24,25^ with 93,766 and 65,903 single cells to compare with our CCC data using the Pearson residuals integration method from Seurat SCTransform integration.

### Differential single-cell RNA-seq analysis

Pairwise differential expression analysis was performed using MAST v1.12.0^76^ on genes expressed in at least 0.25 fractions of the cells. Genes are differentially expressed if a minimum log fold-change threshold is greater than 0.25 and a p-value is less than 0.05.

### Gene set enrichment analysis

The gene set enrichment analysis was performed in clusterProfiler^77^ on the identified differentially expressed genes.

### Trajectory analysis

Using Spearman correlation, similar epithelial cell types across all endometrial conditions and CCC were identified. Then, pseudo-time trajectory analysis was carried out using Slingshot^78,79^ on the PCA projection and CellRank^78^ on the UMAP projection. Diffusion pseudotime was calculated to determine the genes that change across stages of cancer.

### Cell-cell communication analysis

The changes in overall signalling pathway communication between different cell types present within the CCC TME and other endometrial conditions were computed and visualised as dot plots and chord plots using CellChat^80^.

### SNPs detection

The SNPs were detected in the scRNA-seq data using scAllele^81^. The identified SNPs were analysed and visualised using Maftools^82^.

### Drug prediction

Drugs targeting the TME in CCC at different stages of cancer and different cell types were computed using drug2cell^63^ based on the enrichment of drugs for the transcriptional signatures.

## Data Availability

The scRNA-seq data was visualised using an interactive website developed using ShinyCell^83^. The scRNA-seq data generated in this study will be made available on the GEO database. The website files will be made available on GitHub.

## Acknowledgements

We thank Chin Hu Lim for his philanthropic funding support.

